# CRISPR-mediated chromosome deletion facilitates genetic mapping of Vip3Aa resistance gene within complex genomic region in an invasive global pest

**DOI:** 10.1101/2024.07.30.605831

**Authors:** Minghui Jin, Yinxue Shan, Yan Peng, Senlin Chen, Xuanhe Zhou, Kaiyu Liu, Yutao Xiao

## Abstract

Connecting genetic variation to phenotypes and understanding the underlying biological mechanisms has been a fundamental goal of biological genetics. Here, we used the association analysis to identify a Vip3Aa resistance-associated genomic region in a strain of fall armyworm, JC-R, which exhibits >5000-fold resistance to the Bt toxin Vip3Aa. However, through various analytical approaches and fine-scale mapping across different populations, we demonstrated that this genomic region exhibits strong genetic linkage. The chromosome-level genome of JC-R and its parent strain JC-S were assembled, and extensive structural variations in the linkage regions were identified, which could be responsible for maintaining the linkage. To identify the causal variation within this linked region, a chromosome fragment stepwise knockout strategy based on CRISPR/Cas9 was developed. By crossing with the resistant strain and phenotyping segregating offspring on Vip3Aa-containing diet, we identified a chromosomal segment, KO8, containing the resistant gene. Subsequently, we conducted a comprehensive analysis of the variations in the KO8 region using multi-omics approaches, including genomic data, RNA-seq, proteomic, PacBio long read Iso-seq, and phosphoproteomic data. This analysis identified multiple variations in the chitin synthase gene *CHS2*, including amino acid substitution, alternative splicing, and changes in phosphorylation sites. After knocking out the *CHS2*, larvae exhibited over 6777-fold resistance to Vip3Aa. These results demonstrate that the chromosome fragment stepwise knockout strategy is a viable approach for studying complex genomic regions, and highlight the value of comprehensive analysis of genetic variations using multi-omics data. The identified candidate gene could potentially advance monitoring and management of pest resistance to Vip3Aa.

## Introduction

The longest-standing question in genetics is to understand how genetic variation contributes to phenotypic variation^1^. Gene mapping studies aim to establish associations between genomic DNA sequence variants and phenotypic traits^2,3^. Over the past two decades, powerful statistical models have been developed to detect quantitative trait loci and investigate their biological implications across a wide range of phenotypes^3-6^. Genome-wide association studies (GWAS) utilize genotyping arrays to asses genetic variability throughout the genome, serving as a standard platform for testing the association between phenotypes and genetic variants^2^. Single nucleotide polymorphisms (SNPs) employed in GWAS may represent neutral variations that do not affect the traits under study. However, linkage disequilibrium (LD) between SNPs and causal polymorphisms can establish associations with traits. GWAS primarily aim to map genomic regions but typically do not pinpoint causal polymorphisms^7^. The structure of LD is influenced by various factors including recombination, mutation rates, population demographics, and epistatic interactions ^8-10^. However, long-distance LD patterns leads to extensive genomic regions of association where specific variant information is indistinguishable. Tightly linked loci within such long-distance LD regions pose a barrier to resolving loci associated with traits, hindering their segregation. An example is clustered-spikelet rice, an atypical germplasm with the unique trait of multiple complete spikelet. Researchers have endeavored to pinpoint this locus, narrowing it down to a region spanning approximately 22.85-23.85 Mb region on chromosome 6^11-13^. Zhang et al.^14^ devised a strategic cloning approach involving large-scale mutagenesis to identify and characterize loci using mutant lines, revealing multiple structural variations within this long-distance LD region.

The Gram-positive bacterium *Bacillus thuringiensis* (*Bt*) stands as the most economically successful entomopathogen to date, revolutionizing the control of some major pests^15-17^. However, the rapid evolution of pest resistance to Bt toxin poses a serious threat to long-term effectiveness of this technology^18,19^. Better understanding of the genetic basis of resistance to Bt toxin is urgently needed to monitor, delay, and counter pest resistance. Recently, the forward genetic approach of GWAS has been applied to the analysis of Bt resistance mechanisms. For instance, Jin *et al*. conducted a GWAS using a mass backcross population of *Helicoverpa armigera*, pinpointing a 0.67 Mb genomic region associated with Cry1Ac resistance^20^. Guo *et al*. identified a 1.7 Mb region associated with Cry1Ac resistance in *Plutella xylostella* using GWAS^21^. In another study, we utilized 148 individual *Spodoptera frugiperda* of F_2_ progeny generated by DH-R and DH-S, identifying a 0.62 Mb region associated with Vip3Aa resistance through GWAS analysis^22^. These investigations successfully narrowed down the association regions, facilitating the identification of candidate genes through fine mapping. However, when dealing with regions that are excessively long and tightly linked, such as those encompassing complex chromosome structural variations^14^ or chromosome inversions^23,24^, further research is essential to devise strategies for identifying candidate genes without disrupting these tightly linked regions.

In this study, we utilized a forward genetic approach to identify a complex genomic region on chromosome (chr) 18 tightly linked with Vip3Aa resistance in *S. frugiperda* strain JC-R, which shows 5247-fold resistance. Both GWAS analysis of F_2_ progeny and bulked segregation analysis (BSA) of backcross progeny identified a > 5Mb Vip3Aa resistance-associated region on chr18. Fine-scale mapping with another backcross population and a F_6_ population which have constant crossed with wild-type strain narrowed this to a 2.54 Mb region (LOC1). Chromosome-level genome assemblies of JC-R and its parent strain JC-S revealed high structural variation in this region. To overcome linkage challenges in LOC1, we developed a CRISR/Cas9-based method to progressively knockout chromosome fragments. Using this approach, we identified a 0.241 Mb region, KO8, associated with Vip3Aa resistance. Mutil-omics analysis, including sequence comparison, RNA-seq, proteome, Pacbio long read Iso-seq, and phosphoproteomics, between JC-S and JC-R. We found 6 of 14 genes within the KO8 region have amino acid coding variations. chitin synthase 2 (*CHS2*) emerged as a candidate gene with amino acid changes, alternative splicing and phosphorylation modification variations in JC-R. Knockout out *CHS2* in susceptible strain using CRISPR/Cas9 significantly decrease susceptibility to Vip3Aa, confirming that mutation of this gene can cause resistance to Vip3Aa. Our study highlights *CHS2* as a key gene for Vip3Aa resistance and demonstrates a rapid method for identifying trait-controlling genes in complex genomic regions.

## Results

### Selection for resistance to Vip3Aa

We analyzed the Vip3Aa-resistance strain of fall armyworm, JC-R, derived from the JC-S strain originating in Jiang Cheng, Yunnan Province, China, in 2019. After 42 generations of selection with Vip3Aa in the laboratory, the lethal concentration for 50% of larvae (LC_50_) was 5247 times higher for JC-R compared to JC-S (250.35 and 0.048 μg toxin per cm^2^ diet, respectively, Figure 1A, Table S1). Selection for resistance to Vip3Aa in JC-R did not confer cross-resistance to Cry1Ab, Cry1Fa, or Cry2Ab toxins (Table S1). Transgentic Vip3Aa maize demonstrated high efficacy against JC-S but significantly reduced efficacy against JC-R (Figure 1B, Table S2). Bioassay of progeny from reciprocal crosses between JC-S and JC-R indicated that resistance inheritance at a concentration of 0.4 μg Vip3Aa per cm^2^ diet was autosomal and partially recessive (Table S3, mean *h* = 0.075, where *h* = 0 indicates completely recessive and *h* = 1 indicates completely dominant resistance).

**Figure 1.**
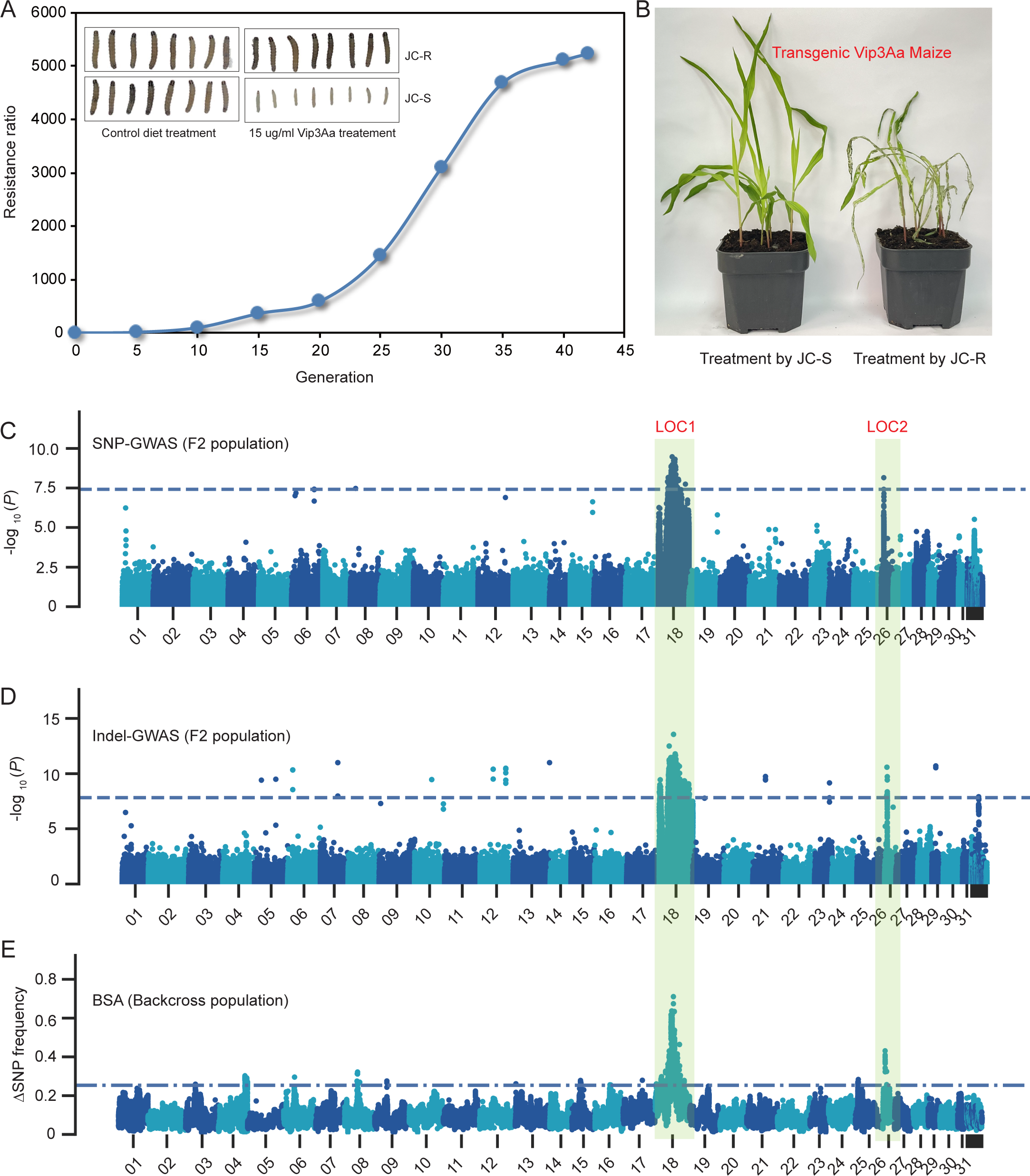
Genomic regions associated with *S. frugiperda* resistance to Vip3Aa. (A) Vip3Aa resistance selection process of JC-R strain. Phenotype of JC-R and its original susceptible JC-S strain feeding on control diet and 15 μg/ml Vip3Aa containing diet. (B) Phenotype of JC-S and JC-R feeding on transgentic Vip3Aa maize. Manhattan plots of F_2_ population: (C) SNPs and (D) indels for all 31 chromosomes. Horizontal dashed lines indicate the threshold for significant association. (E) Manhattan plot for bulked segregant analysis of backcross population. The top 1% Δ(SNP-index) region was identified as candidate region. Common associated regions (LOC1 in chromosome 18 and LOC2 in chromosome 26) of GWAS and BSA were labeled in green.

### Genome-wide association study (GWAS) and bulked segregant analysis (BSA)

We conducted GWAS analysis using SNPs and indels identified from 153 *S. frugiperda* larvae. These larvae were F_2_ progeny generated by a single-pair cross between a JC-R female and a JC-S male followed by mass mating among their F_1_ progeny. Resistance phenotype were scored as larval weight 7d after neonates were placed on diet with 0.4 μg Vip3Aa per cm^2^ diet. Based on analysis of linage disequilibrium (LD) with a threshold of 0.5 for *r*^2^, two regions on chr18 and chr26 were associated with resistance based both on SNPs and indels (Figure 1C, D). LOC1 on chr18 was ranging from 5.21-10.72 Mb based on SNPs, and 4.15-10.85 Mb based on indels. LOC2 on chr26 was ranging from 2.98-3.04 Mb based on SNPs and indels (Figure 1C, D).

LOC1 is too large for effective candidate gene identification, prompting us to conduct additional analysis using Bulk Segregant Analysis (BSA). Genomic DNA from Vip3Aa-resistant (F_2_-R, individuals surviving on 5.0 μg Vip3Aa per cm^2^ diet, n = 48) and Vip3Aa-susceptible (F_2_-S, individuals showing inhibited growth on 0.2 μg Vip3Aa per cm^2^ diet, n = 48) larvae from backcross progeny was used for BSA. Same genomic regions to GWAS which located on chr18 (5.63-10.78 Mb) and chr26 (2.98-3.04 Mb) were identified (Figure 1E). Integrating GWAS (SNPs and indels) with BSA results, we identified an overlapping 5.09 Mb region of LOC1, spanning from 5.63 Mb to 10.72 Mb, for further fine-scale mapping.

### Fine-scale mapping of Vip3Aa resistance

We conducted fine-scale mapping using 96 backcross larvae derived from a single-pair cross between a JC-R female and a JC-S male, followed by mating the single F_1_ male progeny with a JC-R female. Larvae surviving exposure to diet treated with 5.0 μg Vip3Aa per cm^2^ diet were scored as resistant. Our results show a highly significant association between resistance and each of seven SNP markers tested spanning from 5.86 to 10.70 Mb on chromosome 18 (*P* < 10^-15^, Figure 2A, Table S4). Specifically, five markers from 6.93 to 10.46 Mb exhibited complete association with resistance (Figure 2A, Table S4). We hypothesised that this complete association may be attributed to the BC population’s failure to disrupt linkage in the LOC1 region. Subsequently, we established an F_6_ population through through continuous Vip3Aa screening and consistent crosses with wide-type individuals in an effort to break this linkage (Figure S1). Although the F_6_ population narrowed down the candidate regions, a 2.54 Mb segment (ranging from 7.52 to 10.06 Mb) still showed complete association with Vip3Aa resistance (Figure 2A, Table S5).

**Figure 2.**
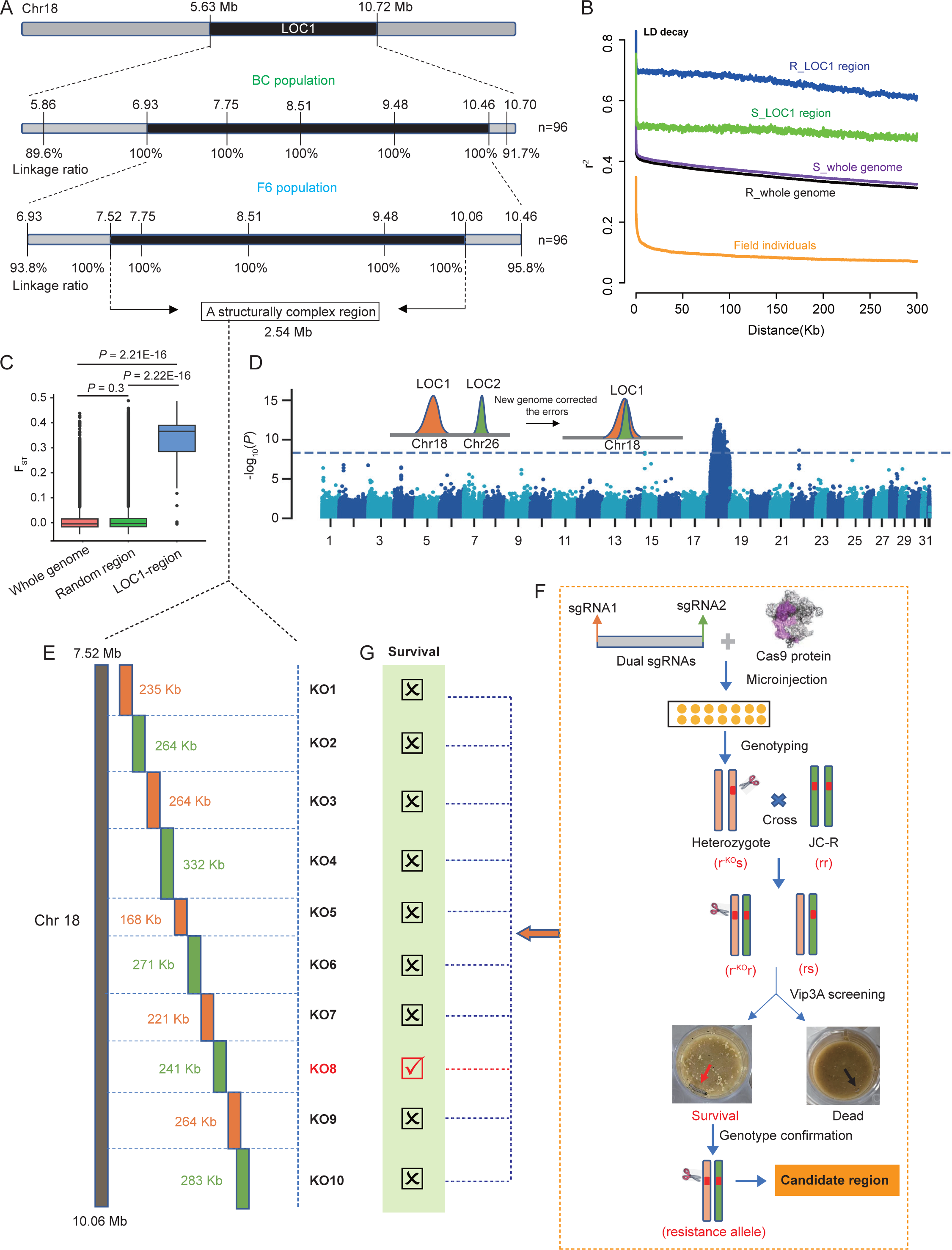
Chromosome deletion via CRISPR facilitate fine mapping of Vip3Aa resistant regions in JC-R. (A) Fine mapping of the LOC1 in chromosome 18. Seven markers were designed to perform the fine mapping using a backcross (BC) population. The frequency of each allele in the resistant individuals (n = 96) were determined. Detail information was listed in Table S4. Another seven markers were designed to perform the fine mapping using a F_6_ population. The frequency of each allele in the resistant individuals (n = 96) were determined. Detail information was listed in Table S5. (B) The linkage disequilibrium (LD) decay in LOC1 region of resistant individuals (R_LOC1), LOC1 region of susceptible individuals (S_LOC1), whole genome of resistant individuals (R_whole genome), whole genome of susceptible individuals (S_whole genome), and field populations. (C) Genetic differentiation (F_ST_) of whole genome, random selected region and LOC1 region between RR individuals and SS individuals (153 F_2_ individuals divided into RR and SS based on phenotype feeding on Vip3Aa containing diet). (D) GWAS re-analysis results based on the new-assembled JC-S genome. The original two LOCs were corrected to a single LOC. (E) Schematic diagram of segmented knockout of chromosome fragments in the narrowed LOC1 (7.52-10.06 Mb) using CRISPR. The length of each chromosome fragments is marked. (F) The work flow of screening homozygous variant alleles through hybridization between chromosome fragment knockout individuals and JC-R resistant individuals. Survivors after Vip3Aa screen contained two resistant allele (r^KO^r, r^KO^ allele obtained from knockout individual, r allele obtained from JC-R individual). (G) Bioassay results showed the progeny of KO8 (ranging from 9.275 to 9.516 Mb) with JC-R have survivors feeding on 5.0 μg Vip3Aa per cm^2^ diet.

We conducted a linkage disquilibrium (LD) decay analysis using the LOC1 region in JC-R and JC-S, as well as whole-genome regions and field populations. The results revealed that the LOC1 region in JC-R exhibited the most pronounced LD decay (Figure 2B). Additionally, we performed fixation index (F_ST_) analysis to assess differentiation within LOC1. The results indicated significant differentiation compared to both randomly selected regions and the whole genome (Figure 2C). The average F_ST_ value for LOC1 was 0.34, markedly higher than 0.012 for random regions and 0.008 for the entire genome, suggesting substantial genetic differentiation within LOC1. These findings collectively indicate that the narrowed LCO1 region is structurally complex.

Despite efforts with the backcross and F_6_ population, genetic linkage in the LOC1 region remained unbroken. To further analyze this complex region, we assembled chromosome-level genome of JC-R and JC-S using combined HiFi reads and Hi-C data, achieving scaffold N50 of 13.35 Mb and 13.32 Mb, respectively (Table S6). Using the new assembled JC-S genome as a reference, we re-analyzed the GWAS data. Interestingly, the original two Vip3Aa resistance associated loci now appeared as a single locus (Figure 2D). Comparison with the genome of ZJ-Sfu^25^ revealed that the ZJ-Sfu genome incorrectly assembled two loci from a single chromosome into separate positions. However, the new genome assembly did not resolve the issue of large resistance linkage regions, with the association region still exceeding 5 Mb (Figure 2D). A genome-wide scan of structural variations in JC-R and JC-S identified chromosome 18 as having highest number of structural variations (Figure S2). Using 1 Mb as sliding window, we found that there were the most structural variations within the resistance linkage region (Figure S3). These findings confirm that the Vip3Aa resistance linkage region on chromosome 18 is structurally complex.

### Rapid screening of chromosome regions controlling Vip3Aa resistance *via* CRISPR mediates chromosome deletion

Due to the difficulty in disrupting the tight linkage within LOC1, we developed an elimination method using CRISR/Cas9 to sequentially knock out chromosome fragments (Figure 2E). The 2.54 Mb Vip3Aa resistance linkage region was divided into ten knockout (KO) regions ranging from 168 kb to 332 kb (Figure S4). Dual sgRNAs were designed flanking the beginning and ending of each KO region. After genotyping, we crossed heterozygous (*r^KO^s*) individuals with JC-R (*rr*) individuals to produce F_1_ progeny (Figure 2F). Subsequently, the F_1_ progeny were screened using a diet containing 5.0 μg Vip3Aa per cm^2^. If the knockout region contained the resistance genes, approximately half of the F_1_ progeny would carry a resistant genotype (*r^KO^r*, knockout allele and resistant allele form homozygouts resistant alleles). Following screening of all KO regions, survivors were only found among the the F_1_ progeny of KO8 (Figure 2G, 40/96, *P* = 0.1, observed genotype ratio not significantly different from the expected 1:1 genotype). This indicates that the Vip3Aa resistance gene in JC-R resides within the KO8 region, spanning from 9.275 Mb to 9.516 Mb. By generating heterozygotes through CRISPR and subsequent backcrossing with resistant individuals, we efficiently identified the region controlling the resistance locus within a single generation. This approach provides an effective solution for tackling complex genomic regions.

### Identification of genes associated with Vip3Aa resistance in JC-R

In the KO8 region, a total of 14 genes were annotated. Amino acid sequence comparison between JC-R and JC-S revealed variations in six genes: protein phosphatase (*phlpp*), hypothetical protein (*HP*), alpha-methyldopa hypersensitive protein (*AMHP*), defensin precursor (*DP*), MPN domain-containing protein (*MPN*) and chitin synthase 2 (*CHS2*) (Figure 3A). Specifically, the *phlpp* gene exhibited a 101 amino acid variation due to a 4004 bp LTR retrotransposon insertion at the transcription start site. The *HP* gene showed a 53 amino acid deletion and 11 amino acid substitutions. The other four genes also exhibited amino acid substitutions. RNA-seq analysis of midguts from fourth instar larvae showed substantial expression (FPKM > 1) of nine out of these 14 genes. However, none of these genes showed significant differences in expression between JC-R and JC-S (Figure 3A). The proteome analysis of midguts from fourth instar larvae revealed substantial protein expression for only three genes, none of which showed significant differences between JC-R and JC-S (Figure 3A). Notably, only CHS2 and CHS1 were found to be expressed in the plasma membrane based on the expected locations of their encoded proteins. PacBio long read Iso-seq analysis of midguts also detected alternative splicing variations in *ypt7* and *CHS2* genes. Additionally, a hosphoproteome analysis of midguts aimed to detect post-translational modification variations associated with Vip3Aa resistance in JC-R. We identified 2 phosphorylation sites (PS) in *Nelf*, 2 PSs in *ypt7*, 3 PSs in *MPN* and 14 PSs in *CHS2*. Among these phosphorylation sites, only 10 out of 14 PSs in the *CHS2* gene showed significant differences between JC-R and JC-S (Figure S5).

**Figure 3.**
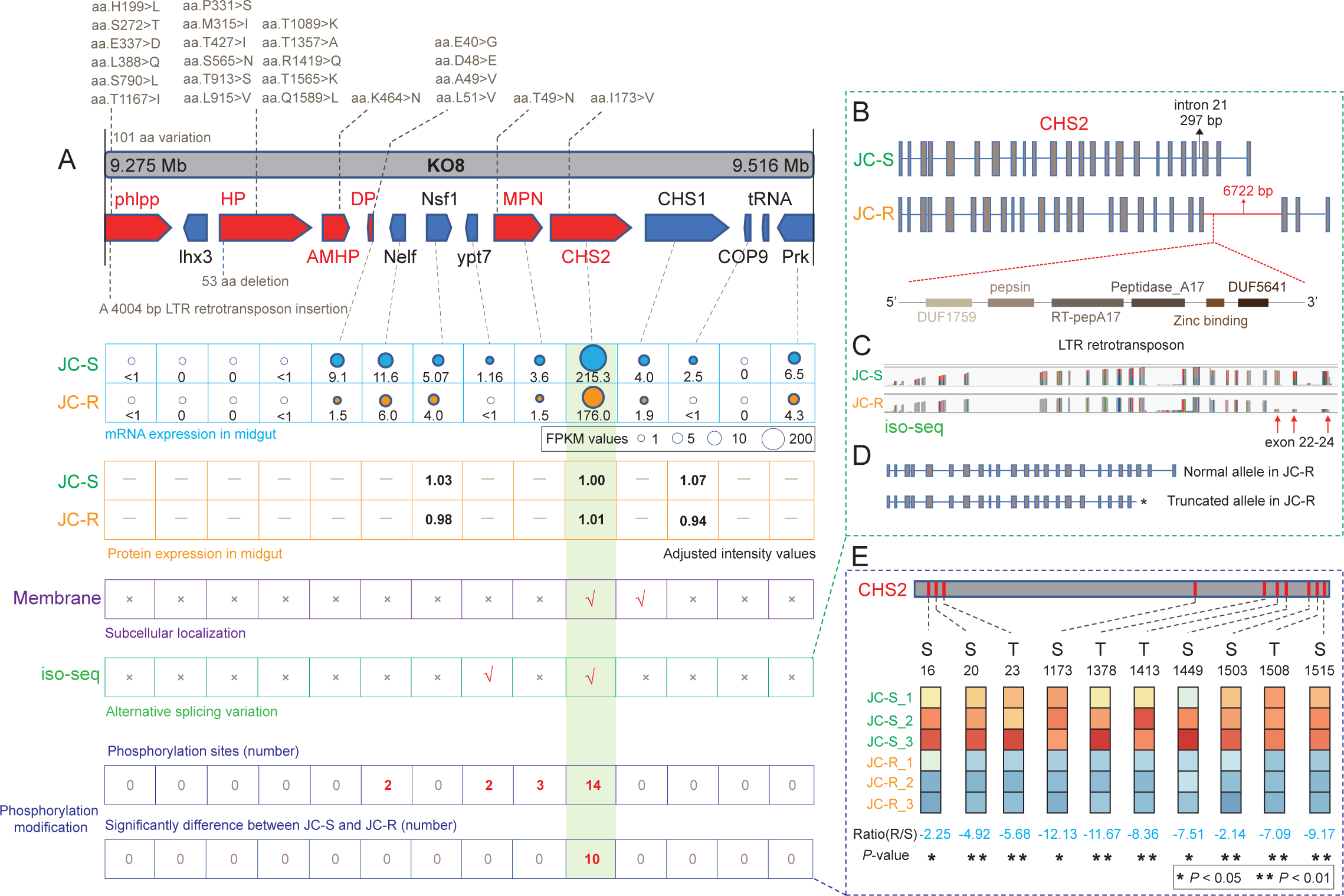
Multi-omics data identify variations in Vip3Aa resistance-related genes. (A) Schematic diagram of annotated genes and it’s mutation information between JC-S and JC-R within KO8 region. 1) Fist layer - amino acid variation: genes have amino acid variation between JC-S and JC-R have been marked in red. Amino acid substitution and other mutation types of amino acid were labeled in gray; 2) Second layer - mRNA expression in midgut: gene expression data obtained from RNA-seq of midgut from JC-S and JC-R. The FPKM values were labeled. If the gene’ FPKM <1 or =0, considered it did not expressed in midgut; 3) Third layer - protein expression in midgut: protein expression data obtained from proteome of midgut from JC-S and JC-R. The numerical value represents adjusted intensity values; 4) Fourth layer - subcellular localization: The prediction of subcellular localization were performed in DeepTMHMM (https://dtu.biolib.com/DeepTMHMM). Check marks indicates it located in membrane; 5) Fifth layer - alternative splicing variation: isoform-sequencing (full-length transcriptome) of midgut from JC-S and JC-R was performed. Check marks indicates these genes have alternative splicing variation between JC-S and JC-R; 6) Sixth layer - protein post-translational modifications: the phosphorylated proteome of midgut from JC-S and JC-R was sequenced. The numerical value represents the number of identified phosphorylation sites and significantly difference modified phosphorylation sites between JC-S and JC-R. (B) A LTR retrotransposon insertion was identified in the intron 21 of *CHS2* in JC-R. Gene structure of *CHS2* in JC-S and JC-R and the domains in the LTR retrotransposon were showed. (C) Expression of *CHS2* visualisation results based on iso-seq data. (D) The insertion of LTR retrotransposon introduced a premature termination codon, resulting in a deletion of exon 22-24. (E) Ten significantly different phosphorylation sites in CHS2 protein between JC-S and JC-R. Blue represent the reduced modification of phosphorylation sites in JC-R.

Based on multi-omics data, we identified *CHS2* as a candidate resistance gene against Vip3Aa. The *CHS2 gene exhibits several key characteristics: it harbors amino acid substitutions, shows mRNA and protein expression in the midgut, it localized in the membrane, displays alternative splicing variations, and exhibits significantly differences in* phosphorylation sites between JC-R and JC-S (Figure 3A). Detailed analysis of alternative splicing in *CHS2* revealed a 6425 bp LTR retrotransposon inserted in intron 21 (Figure 3B). This insertion introduced a premature termination codon, resulting the deletion of exon 22-24 (Figure 3C, 3D). Interestingly, both the variant and normal alleles are expressed in JC-R individuals, as visualized in our Iso-seq results (Figure 3C). Consequently, conventional analysis methods did not detect significant differences in mRNA and protein expressions of *CHS2* between JC-R and JC-S. Furthermore, analysis of phosphorylation sites in *CHS2* showed that ten sites were significantly reduced in JC-R, ranging from a 2.14-fold to 12.13-fold decrease (Figure 3E).

### Knockout CHS2 decreases susceptibility to Vip3Aa

We employed the CRISPR/Cas9 gene editing system to investigate the impact of disrupting the *CHS2* gene by introducing a premature stop codon on susceptibility to Vip3Aa. Through the use of dual sgRNAs, we generated a strain that is homozygous for a 223-bp deletion in *CHS2* (Figure 4A, 4B). Bioassay results clearly demonstrate that the *CHS2* knockout strain (CHS2-KO) confers > 6777-fold resistance to Vip3Aa (Figure 4C). Even at the highest tested concentration (250 μg Vip3Aa per cm^2^ diet) of Vip3Aa, larvae from the CHS2-KO strain exhibited a remarkably high level of resistance, with larval mortality only reaching 4.2%. Fluorescence in situ hybridization (FISH) results confirmed the expression of *CHS2* mRNA in the midgut brush border and epithelium cells (Figure 4D). Conversely, in the CHS2-KO strain, *CHS2* mRNA was not detectable in the midgut brush border or epithelium cells, instead, it was enriched in the outermost layer of the midgut basal lamina (Figure 4E). Further investigation is needed to determine whether the mislocalization of *CHS2* mRNA residues to the basal lamina or the potential overexpression of the homologous gene *CHS1*, as a compensatory response to *CHS2* deletion. However, our findings clearly indicate that after *CHS2* gene knockout, midgut cells no longer express intact *CHS2* mRNA (Figure 4E). Additionally, upon *CHS2* gene knockout, we observed the disappearance of the peritrophic membrane (PM) (Figure S6). These observations led us to speculate on the mechanism through which Vip3Aa exerts its toxicological effects. Specifically, the Vip3Aa protoxin enters the midgut and initially binds to the PM, and subsequently to midgut receptors, either as single Vip3Aa toxins or polymers enriched after PM binding. This interaction leads to midgut perforation or the induction of apoptosis or autophagy within cells (Figure 4F). In the Vip3Aa resistant JC-R strain, the function of the *CHS2* gene is influenced by several factors including amino acid substitution, alternative splicing variation, and phosphorylation variation. These variations in the *CHS2* gene may effect the synthesis of the PM or the binding of Vip3Aa to the midgut, thereby contributing to the high-level resistance in JC-R (Figure 4G). After knockout the *CHS2* gene, insects are unable to form a PM. This may prevent the enrichment of Vip3Aa toxin by the PM or its binding to the midgut, resulting in completer resistance (Figure 4H).

**Figure 4.**
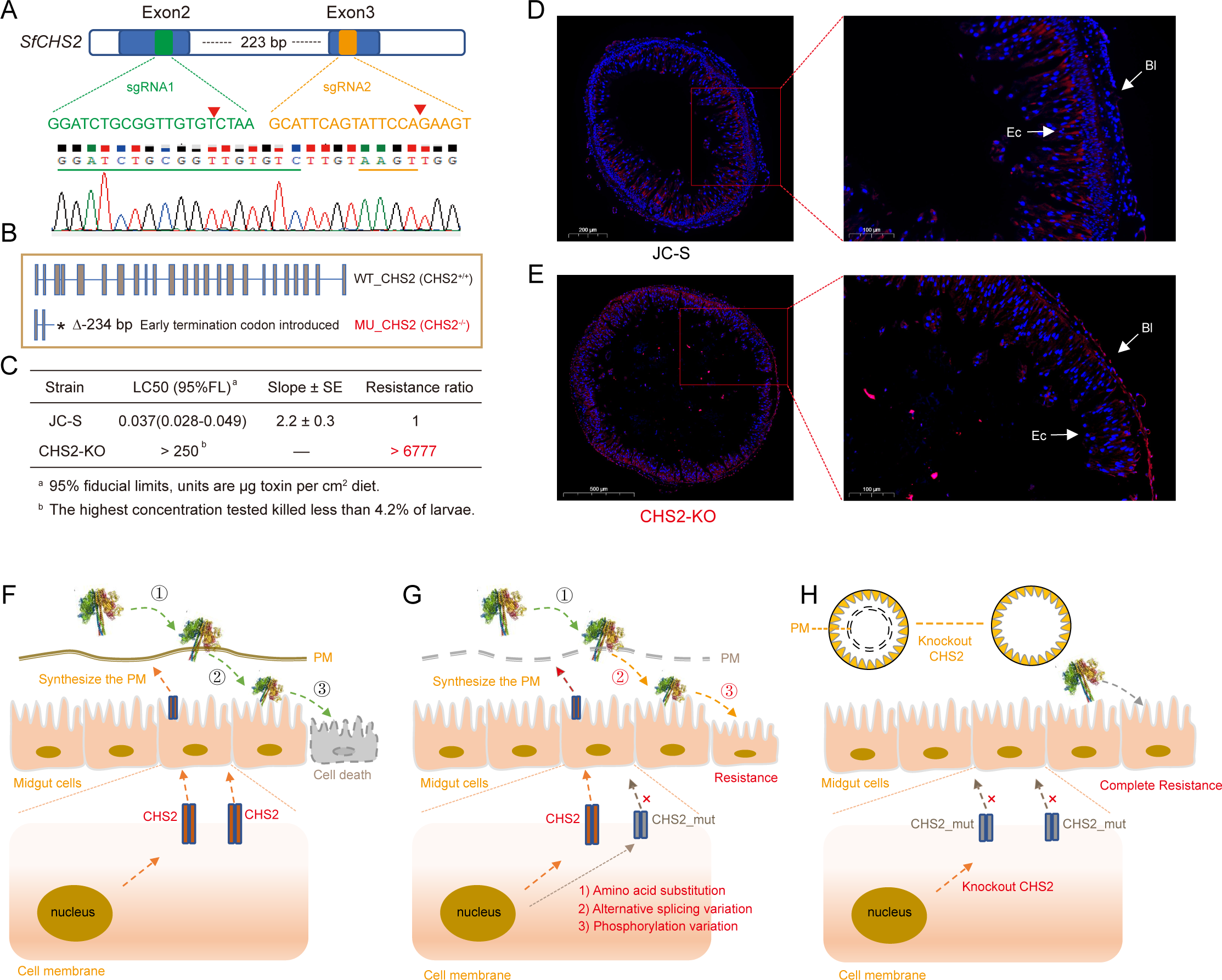
Validation of CHS2 function via CRISPR/Cas9. (A) CRISPR/Cas9-mediated knockout of CHS2 gene using dual sgRNA. A 223-bp deletion mutation type selected for homozygous screen. (B) The 223-bp deletion introduced a premature termination codon. (C) Bioassay results of CHS2-KO strain exposed to Vip3Aa protoxin. The CHS2-KO strain exhibited a high level of resistance to Vip3Aa, even at the highest concentration (250 μg Vip3Aa per cm^2^ diet) of Vip3Aa, with larval mortality only reaching 4.2 %. (D) Fluorescence in situ hybridization (FISH) results of CHS2 mRNA in the midgut of JC-S strain. Red fluorescence represents the expression of CHS2 mRNA. Ec, epithelium cells; Bl, brush border. (E) FISH results of CHS2 mRNA in the midgut of CHS2-KO strain. (F) Proposed primary model of the insecticidal process of Vip3Aa. 1: Vip3Aa in the midgut can bind to peritrophic matrix (PM) through domain V of Vip3Aa^62^; 2: single Vip3Aa or polymer after PM enrichment passing the PM, interacting with the potential receptors on the midgut; 3: eventually it will lead to cell death forming pores on the midgut cells. (G) Proposed primary model of Vip3Aa resistance mechanism in JC-R. Muti-type of mutations in the *CHS2* gene may effect the synthesis of PM (2) or the binding of Vip3Aa to the midgut (3), causing high-level resistance to Vip3Aa. (H) Knockout of *CHS2* gene results in an inability to synthesis the PM, which in turn affects the binding of Vip3Aa to PM or the binding to *CHS2* in the midgut, ultimately leading to complete Vip3Aa resistance.

## Discussion

GWAS utilize genotyping arrays to assess genetic variation across the genome, serving as the standard method to investigate the correlation between a phenotype and genetic variants^3,26^. However, the trait-associated region measure by GWAS still has limitations. Many genetic variants identified through GWAS may have negligible effects on the phenotype, and including numerous such variants reduces statistical power^27^. Furthermore, structural variations, which are not typically directly genotyped and may not be accurately imputed, can result in large linkage disequilibrium regions that diminish the ability to detect associations^28,29^. The precise localization of traits within complex genomic regions thus remains a significant challenge. In this study, we identified a genomic region characterized by a complex gene structure that shows a strong association with Vip3Aa resistance through association analysis including GWAS and BSA analysis.

Despite conducting fine-scale mapping using a backcross population and an F_6_ population, we encountered challenges in breaking the linkage within a complex genomic region. To address this issue, we adopted an innovative approach based on stepwise chromosome fragment knockout using CRISPR/Cas9 technology. Through this method, we successfully narrowed down the complex genomic region from 3.5 Mb to 0.24 Mb. Complex chromosome structural variation around the casual locus does make it difficult to fine-scale mapping. This challenge is exemplified by previous studies on the clustered-spikelet locus in rice, where despite various strategies, the identified linkage regions remained large and difficult to refine^11-13^. Notably, one study achieved success by analyzing a large number of mutagenized individuals, leading to the identification of a mutant that finally pinpointed the causal gene within the complex region^14^. Chromosome inversions represent another type of structural variation that complicates recombination processes^30,31^. When inversions are associated with complex phenotypes, identifying causal mutations becomes particularly challenging because mutations within the inversion region often exhibit strong associations with phenotype^24,32,33^. Our findings underscore the efficacy of the chromosome fragment stepwise knockout strategy, especially in species with shorter generation times like insects. This approach proves both time-efficient and effective for pinpointing causal genes within complex genetic regions.

CRISPR/Cas9 system as a powerful genome editing tool have revolutionized functional genomics studies in insect pests^34,35^. It have been widely used in insects to introduce a frameshift mutations^21,36-38^, single nucleotide substitutions^20^, and chromosomal deletions^39-42^ in target genes. With the dual sgRNAs strategy, the chromosome fragments deletion can be visualized by electrophoretic bands. This methodology has been successfully applied to demonstrate chromosome deletions in species such as *H. armigera*^39-41,43^, and *S. frugiperda*^42^. In this study, we adapted this chromosomal fragments knockout approach to fine-map complex genetic regions. By designing overlapping sgRNAs, we divided the complex LOC1 region into 10 segments for chromosomal knockout, successfully generating knockout heterozygotes for each segment. We have conceived a hybridization strategy: based on the premise of a 1:1 genotype separation ratio (S^KO^R:SR =1:1) between knockout heterozygotes (S^KO^S) and resistance individuals (recessive inheritance, RR); the offspring were screened using a lethal concentration of Vip3Aa; if the region containing resistance gene, the survival rate of the offspring should be 50%. Through this strategy, we identified the KO8 region containing the candidate genes associated with resistance. This strategy enabled rapid and efficient screening of key genomic regions controlling traits within a single generation, facilitating the precise fine-mapping of complex genetic regions. By leveraging CRISPR/Cas9 technology in this manner, researchers can expedite the identification of genetic determinants underlying important traits in insect populations.

The assembly of a high-quality reference genome serves as the foundation for accurate gene identification and genetic variation analysis^44^. In this study, we initially utilized the published reference genome of *S. frugiperda* for conducting GWAS analysis, which led to the identification of two loci on two different chromosomes^25^. To further analyze the genomic structure of the complex linked region LOC1, we generated two chromosome-level reference genomes for JC-S and JC-R. Using the updated reference genome of JC-S, we re-performed variant detection and GWAS analysis, which identified only one locus on chromosome 18 associated with Vip3Aa resistance. Upon sequence alignment, it was discovered that a region originally belonging to chromosome 18 had been incorrectly assembled into chromosome 26 in the previous genome version, resulting in the identification of two separate loci related to Vip3Aa resistance. Sequence analysis revealed the presence of numerous structural variations and repetitive sequences in this complex genomic region, which likely contributed to assembly errors in previously genome version^45,46^. These findings underscore the critical importance of having a high-quality reference genome for accurate gene mapping and genomic analysis. A precise genome assembly is essential for correctly identifying genetic loci associated with traits of interest, particularly in regions characterized by complex genomic structures and variations.

Evolution of Bt resistance is the most relevant threat to sustainability of Bt insecticidal proteins. Understanding the genetic basis of resistance to Bt toxin is crucial for effective resistance monitoring and management practices. Several receptors for Cry toxins, such as cadherin (CAD)^47^ and ABC (ATP-binding cassette) transporters^48^ have been identified in insects. Resistance to Bt toxins often arises from mutations affecting these receptors. Various types of mutation in CAD have been associated with Cry resistance, including fragment deletion^49^, amino acids mutation^47,49-52^, single point mutation causing mislocalization^53^, reduced expression^54^, and alternative splicing^55^. Similarly, mutations in ABCC transporters have been linked to Cry toxin resistance, encompassing amino acids changes^56^, single point substitution^57^, and reduced expression^37^. In some cases, resistant pest populations exhibit multiple variants of these receptors. For example, in the cotton bollworm, *H. armigera*, mutations affecting both major receptors for Cry toxins - the midgut cadherin and ABC transporter - have been identified within a single strain resistant to Cry toxins^37^. This highlights the complexity of resistance mechanisms and the potential for pests to evolve resistance through mutations affecting multiple receptor types simultaneously.

In this study, we conducted a thorough analysis of variations within a candidate region using a multi-omics approach that integrated genome sequencing, RNA-seq, proteomics, PacBio long read Iso-seq, and phosphoproteomics. This comprehensive strategy enabled us to identify numerous variants within the candidate gene *CHS2*. The use of a single omics analysis alone would have been insufficient for a comprehensive examination of gene variations. For instance, within the KO8 region, we identified 11 genes, among which six exhibited amino acid variations, and two transposon insertions were also detected. Moreover, relying solely on gene expression data to identify potential candidate genes proved challenging, as no genes showed differential expression between the JC-R and JC-S strains. Additionally, two genes within this region exhibited alternative splicing patterns. Thus, the evaluation of candidate gene variations using a single omics approach is inherently limited. By employing a multi-omics strategy, we were able to capture a more complete picture of genetic variations, alternative splicing events, and post-translational modifications that may contribute to complex phenotypes such as resistance to insecticidal proteins like Vip3Aa. This integrative approach is crucial for unraveling the intricate molecular mechanisms underlying trait variation in insect pests and other biological systems. Liu et al.^58^ identified a alternative splicing in *CHS2* gene was related to Vip3Aa resistance in a lab-selected *S. frugiperda* Vip3Aa resistance strain. In this study, we also identified the same alternative splicing variation in *CHS2*. Additionally, we identified a novel amino acid substitution, I173>V, in the JC-R strain that was not previously reported. Furthermore, our investigation revealed 14 phosphorylation sites in *CHS2*, with ten of these sites showing significantly reduced phosphorylation levels in the JC-R strain compared to JC-S. Protein phosphorylation, one of the main protein post-translational modifications, is required for regulating various life activities^59-61^. Protein phosphorylation is involved in various important biological activities, such as protein subcellular localization, interaction, stabilization and activation^59^. Variations in protein phosphorylation modification levels could affect the function of proteins.

Knocking out *CHS2* gene, the larvae showed extremely high resistance to Vip3Aa, confirming the pivotal role of this gene in mediating Vip3Aa resistance. However, the knockout resulted in a premature termination and loss function of the entire gene, which only proves that this gene has a key role in the Vip3Aa resistance. In our resistance JC-R strain, *CHS2* gene exhibits alternative splicing, amino acid substitution and reduced phosphorylation levels. However, the specific role of each of these variants in conferring resistance, and whether one variant predominates as the main effector, requires further investigation. Recently, Jiang et al.^62^ demonstrated that Vip3Aa domins Ⅱ Ⅲ could bind the midgut epithelium, while domain Ⅴ can bind the peritrophic matrix via its glycan-binding activity, contributing to Vip3Aa insecticidal activity. Similar to the previous study, we found the peritrophic matrix completely invisible after knockout the *CHS2* gene. In the midgut of JC-S, we confirmed that the *CHS2* mRNA expressed in the midgut brush border and epithelium cells. In the CHS2-KO strain, midgut cell no longer express the intact *CHS2* mRNA, whereas, it can be detected in the outermost layer of midgut basal lamina. Whether the residues of *CHS2* mRNA are mislocalised to the basal lamina, or the homologous gene *CHS1* showed overexpression due to the compensatory effect of *CHS2* deletion still need further study.

In conclusion, our study demonstrates the effectiveness of employing CRISPR/Cas9-based chromosomal fragment deletion to aid in fine-scale mapping of complex genomic regions. By integrating gene editing with hybridization strategies, we successfully identified key regions that control traits within just 1-2 generations. Utilizing a multi-omics approach, we conducted a comprehensive analysis of genetic variations within the identified region and pinpointed the key resistance gene in the laboratory-selected Vip3Aa resistance strain JC-R. Our methodology not only serves as a valuable reference for analyzing complex genetic regions but also sheds light on the mechanisms underlying Vip3Aa resistance.

## Materials and methods

### Insect strains

To start the susceptible strain JC-S of *S. frugiperda*, approximately 150 larvae were collected from more than 10 non-Bt corns fields from Jiang Cheng, Yunnan Province, China, in February 2019. In subsequent generations, JC-S was reared on wheat germ diet in the laboratory without exposure to Bt toxins or other insecticide. In April 2019, we started the Vip3Aa-resistance strain JC-R by exposing approximately 5,000 first instar larvae from JC-S to wheat germ diet treated by Vip3Aa and rearing ∼500 larvae to third instar to continue the strain. JC-R strain had been selected for more than 42 generations with increasing concentrations of Vip3Aa. For maintenance and diet bioassays, larvae were reared on artificial diet at 27± 2 ℃ and 75 ± 10% relative humidity with 14 h light:10 h dark. Adults were provided access to a 10% sucrose solution for nutrition.

### Bt toxins and diet bioassays

Vip3Aa protoxin was provided by the Institute of Plant Protection, Chinese Academy of Agricultural Sciences, Beijing. We used standard diet overlay bioassay to test first or second instar larvae in all experiments ^22,38^. Following the method, larvae were considered dead if they died or weight less than 5 mg after 7d. Similar to previous study, we tested first instar larvae to determine LC_50_, whereas second second instar larvae were used for fine-scale mapping and genetic linkage experiments ^22^. We used second rather first instar larvae in these experiments to increase survival and the size of survivors, which made it easier to extract DNA for subsequent analysis. The LC_50_ and their 95% fiducial limits were calculated with probit analysis using SPSS. Two LC_50_ values were considered significantly different if their 95% fiducial limits did not overlap.

### Transgentic Vip3Aa maize treatment

The seeds of transgentic Vip3A maize and non-Bt corn (Nonghua106) were obtained from the DBN Group (Beijing). Corn was grown in a green-house at about 26 ℃. We transferred first instar larvae to corn when plants were about 30 cm high and had four leaves and one shoot. Ten first instar larvae of JC-S and JC-R were transferred to Vip3a-expressing corn and non-Bt corn, respectively. After 7d, we recorded the growth of each type corn.

### Inheritance of Vip3Aa resistance

To evaluate the inheritance of Vip3Aa resistance in strain JC-R, we allowed 30 virgin male moths from JC-R to mate with 30 virgin moths of the susceptible strain JC-S. The reciprocal cross was also performed. The susceptibility to Vip3Aa toxins of both strain and their F_1_ progeny was tested at 0.4 μg Vip3Aa per cm^2^ diet. We used Fisher’s exact test to determine whether survival differed significantly between the progeny from the two reciprocal crosses. We calculated the dominance parameter *h* as: (Survival of F_1_—Survival of JC-S)/(Survival of JC-R—Survival of DH-S) ^22^.

### GWAS and BSA analysis

#### F_2_ population construction

We conducted a single-pair cross between a JC-S male and JC-R female, then a mass cross with the F_1_ progeny (30 males and 30 females) to obtain F_2_ progeny.

#### Phenotyping by bioassay

To record the phenotypes associated with the Vip3Aa resistance traits, two hundred larvae from the F_2_ generation were reared on a diet treated with 0.4 μg Vip3Aa per cm^2^ diet. The weight of each larva was weighed after feeding on the Vip3Aa diet 7 days.

#### Backcross (BC) population construction

Single-pair crosses between JC-S and JC-R virgin adults to produce F_1_ generation. A male F_1_ was backcrossed to a female JC-R to produce the backcross families. Progeny from the BC family were exposed to two different concentration of Vip3Aa diet. Two hundred larvae feeding on high concentration Vip3Aa diet (5.0 μg Vip3Aa per cm^2^) and then 48 survivors were selected as the F_2_-R group. Another two hundred larvae feeding on low concentration Vip3Aa diet (0.2 μg Vip3Aa per cm^2^) and then 48 individual larvae showed inhibition of growth were selected as the F_2_-S group. These selected larvae were reared to fourth instar on normal diet for DNA extraction.

#### DNA extraction

Genomic DNA was extracted from whole larvae using an EasyPure Genomic DNA kit (TransGen, Beijing, China) according to the manufacture’s recommendations.

#### Genomic sequencing

For GWAS, a total of 153 individual were used for sequencing. Libraries of PE350 were constructed, and sequenced on the BGISEQ-500 platform at BGI-Shenzhen, China. All samples were sequenced with ∼ 20 × depth.

For BSA, each individuals of in pool_F_2_-R, pool_F_2_-S and their parents was sequenced on the BGISEQ-500 platform at BGI-Shenzhen with PE350 libraries. All samples were sequenced with ∼ 20 × depth.

### Variant calling, GWAS and BSA analysis

Raw reads were preprocessed for quality control and filtered using SOApnuke (v1.5.6). Adapters were also trimmed. Clean reads were mapped to the reference genome (ZJ version^63^) using the BWA (v.0.7.17) with the default parameters. Reads sorted were performed using SAMtools (v1.7) and duplicate reads were removed using Picard (v2.17.0). Variant calling was carried out following the Genome Analysis Toolkit (GATK, v4.2.3). The variants for every samples were identified with the HaplotypeCaller module to obtain the genomic variant call format (GVCF) files. Then, the GVCF files were consolidated into a single GVCF file. The SNPs and indels were further filtered using the following criteria: “QD <2.0||MQ<40.0||FS>60.0||MQRankSum <-12.5||ReadPosRankSum <-8.0” for SNPs and “QD < 2.0||FS >200.0||SOR>10.0||InbreedingCoeff <-0.8|| ReadPosRankSum <-20.0” for indels. The SNPs were annotated using SnpEff (v4.3).

Phenotypic data consisted of the weight of each larva after 7d on diet treated with Vip3Aa. GWAS was performed on the SNP (or indel) set using the linear mixed model (LMM) in the program genome-wide efficient mixed model association (GEMMA) (v0.98.1). A Wald test was used to calculate the significance (-lmm1) with the thresholds of 0.05 divided by the number of independent SNPs (or indels). For BSA, we calculated the mean frequency deviation values in two groups using 10-kb sliding windows with less than 50 informative SNPs were discarded to minimize noise. The top 1% values were chosen as the threshold.

### Fine-scale mapping of Vip3Aa resistance

For fine-scale mapping of resistance, we generated a second backcross population. As described above, we started with a cross between a female from JC-R and a male from JC-S to produce F_1_ progeny. Then, a male F_1_ was crossed with a female from JC-R to obtain the backcross population. The larva of BC population were fed with a diagnostic concentration of Vip3Aa (5.0 μg Vip3Aa per cm^2^ diet). 96 survivors were selected for further use. Based on the SNPs calculated from the genomic resequencing data and the reference genome data, we designed primers specific for the exons of functional genes spanning 5.86 to 10.70 Mb on chromosome 18 (Table S7). PCR amplification and sequencing were performed for parent moths. SNPs that were homozygous in JC-S, different homozygous SNPs in JC-R, and heterozygous in F_1_ were selected as informative markers. To evaluate genetic linkage with resistance, the chi-square test for significant deviation between the observed and expected genotype frequencies (*rr*: *rs* = 1:1) was performed.

To narrow the resistance linkage region, a F_6_ population was constructed. We started with a cross between a female from JC-R and a male from JC-S to produce F_1_ progeny. The F_2_ segregating population was obtained by crossing F_1_ population. The progeny of F_2_ population were selected on Vip3Aa diet (5.0 μg Vip3Aa per cm^2^ diet). Male survivors were selected to cross with JC-S female to produce F_3_ population. As in the F_1_ to F_3_ generations, JC-S genotype was introduced and screened with Vip3Aa toxin in the F_4_ to F_6_ generation. 96 survivors of F_6_ were selected for further use. As described above, another seven markers were designed (Table S7). The chi-square test for significant deviation between the observed and expected genotype frequencies (*rr*: *rs*: *ss* = 1:2:1) was performed.

### Linkage disequilibrium (LD) decay and F_ST_ analysis

To estimate and compare the patterns of LD between LOC1 region and other regions in genome, the squared correlation coefficient (r2) between SNPs was analyzed using the PopLDdecay package^64^. The data used to analysis the LD of field individuals was obtained from our previously project ^63^. The genetic differentiation of LOC1 and other genomic regions between JC-S and JC-R was calculated using pixy (v1.2.6)^65^.

### Genome sequencing and assembly of JC-S and JC-R

Genomic DNA of one male pupa of JC-S and JC-R was extracted, respectively. A total of 30 G (75×) and 26 G (68×) HiFi reads of JC-R and JC-S was generated using the Pacific Biosciences Sequel platform. For the construction of Hi-C libraries, genomic DNA of the same pupa was used to construct the Hi-C library using the restriction enzyme Mbol and sequenced *via* Illumina Novaseq. A total of 66.21 G (174×) and 59.23 G (148×) Hi-C data of JC-R and JC-S was obtained. After remove the adapter sequences, the clean PacBio reads were assembled using Hicanu (v2.0)^66^ with default parameters. Hi-C reads were used to map the draft genome with Juicer (v1.5)^67^ and correct misjoins, order and orientation with 3D-DNA (v.180922)^68^. The assembly completeness was evaluated using Benchmarking Universal Singel-Copy Orthologs (BUSCO, v5.1.2)^69^.

### GWAS re-analysis

Reads mapping and SNP calling was performed using the new assembled JC-S genome as reference. A total of 10,772,311 SNPs was calculated and used to GWAS analysis.

### Structural variation identification

Genome assemblies of JC-S and JC-R were aligned using SyRI (v1.0)^70^ with default parameters to identified structural variations between them.

### Chromosome deletion *via* CRISPR/Cas9

We divided the structurally complex region ranging from 7.52 to 10.06 Mb into ten small regions, named KO1 to KO10.

#### Design and synthesis of sgRNA

The sgRNA was designed using the sgRNAcas9 design tool. For each KO region, a pair of sgRNAs was designed at the initial position (forward_sgRNA) and termination position (reverse_sgRNA) (Table S7). The reverse_sgRNA designed in the termination position of KO1, used as the forward_sgRNA of KO2. The same goes for the design of the follow-up sgRNAs. The selected sgRNA target sequence was checked in a search of the *S. frugiperda* genome (JC-S) and GenBank database using sgRNAcas9 design tool, and no potential off-target sites were found. We used the GeneArt Precision gRNA Synthesis Kit (Thermo Fisher Scientific, MA, USA) to synthesize the sgRNA. The *in vitro* transcription was also performed with the same kit according to the manufacturer’s instructions.

### Embryo collection and microinjection

Freshly laid eggs (within 2 h after oviposition) of JC-S were washed from gauze using 1% (v/v) sodium hypochlorite solution and rinsed with distilled water. The eggs were stuck on a microscope slide with double-sided adhesive tape. About 1.5 nL of a mixture of two sgRNAs (150 ng/μL) and Cas9 protein (50 ng/μL) was injected into each egg using Nanoject III (Drummond, Broomall, USA). Cas9 protein (GeneArt Platinum Cas9 Nuclease) was purchased from Thermo Fisher Scientific (Shanghai, China).

### Mutagenesis detection and Vip3Aa screening

Genomic DNA from exuviates, larvae and adults were extracted using a Multisource Genomic DNA Miniprep Kit (Axygen, New York, USA) according to the manufacturer’s recommendations. Primers were designed flanking the CRISPR target sites and used for PCR amplification (Table S7). A smaller PCR fragment is expected to be amplified in the mutated strains compared with wild type is a fragment deletion occurred. After we obtained the heterozygote (*r^KO^s*), we cross it with JC-R (*rr*) to produce F_1_ progeny (*r^KO^r*, *rs*). The F_1_ progeny were feed on 5.0 μg Vip3Aa per cm^2^ diet to detect if survivors can be found. The genotype of survivor was determined using their exuviates.

### Multi-omics identified the candidate gene associated with Vip3Aa resistance

#### RNA-seq of midguts from JC-S and JC-R

From midguts of fourth instar larvae of JC-S and JC-R reared on untreated diet, we extracted total RNA using the TRIzol reagent kit. After quality assessment, six (each strain with three biological replicates) sequencing libraries were constructed and then sequenced on the HiSeq platform. After removing reads containing adapters, low-quality reads, etc, the clean reads were obtained. HISAT (v2.2.4) was used to map the clean data to the reference genome (JC-S). The expression level of genes were estimated by RSEM and normalized using the fragments per kilobase of transcripts per million mapped reads (FPKM). Differentially expressed genes were identified as (|log2 (fold-change)| ≥ 1.5 with q < 0.05) using the DESeq2 (v3.16).

### Proteome analysis of midguts from JC-S and JC-R

Total protein of midguts were extracted using the Tissue Protein Extraction Kit. After trypsin digestion, the resultant peptides were dried using vacuum centrifugation. The separated peptides were analyzed in Orbitrap Exploris 480 with a nano-electrospray ion source. The detail analysis of nano-HPLC-MS/MS were similar to previous reports^71^. Raw data were processed and analyzed via Spectronaut X, with default parameters. We set 1% as the q-value (FDR) cutoff for precursors and protein levels. Decoy generation was set to mutated, which is similar to scrambled, but only applies a random number of AA position swaps (min = 2, max = length/2). All selected precursors passing the filters were used for quantification. After Student’s *t*-Test, 1.2-fold change and *p*-value < 0.05 was set as the threshold to screen differentially expressed proteins.

### Subcellular localization prediction

The prediction of subcellular localization were performed in DeepTMHMM (https://dtu.biolib.com/DeepTMHMM).

### Pacbio long read Iso-seq analysis of midguts from JC-S and JC-R

Total RNA was isolated using the TRIzol reagent kit. The RNA sample from 6 individuals of JC-S and 6 individuals from JC-R were pooled to construct the cDNA library using the SMARTer PCR, respectively. Next, the cDNA library was size-fractioned to 0.5–1, 1–2, 2–3 Kb, and >3 Kb and single-molecule sequencing on Pacbio RS II was performed using P6-C4 kit with 240 min of movie time for each flow cell, respectively (Nextomics Biosciences Institute, Wuhan, China). The Pacbio raw reads were classified into Circular Consensus Sequences (CCS) and non-CCS subreads by searching for the presence of adapters. The long subreads were aligned to the genome using GMAP^72^.

### Phosphoproteome analysis

Total protein of midguts were extracted using the Tissue Protein Extraction Kit. The concentration of protein was determined with BCA kit. After trypsin digestion, the peptide mixtures were incubated with IMAC microspheres suspension with vibration. To elute the enriched phosphopeptides, the elution buffer containing 10% NH_4_OH was added and the enriched phosphopeptides were eluted with vibration. The supernatant containing phosphopeptides was collected and lyophilized for LC-MS/MS analysis. Peptides were separated and were analyzed in Orbitrap Exploris 480 with a nano-electrospray ion source. The MS/MS data were processed using Proteome Discover search engine (v2.4) and then searched against the reference genome. Oxidation on Met, acetylation on protein N-terminal, met-loss on Met, met-loss+acetyl on Met and phosphorylation(S/T) were specified as variable modification. After Student’s *t*-Test, 1.2-fold change and *p*-value < 0.05 was set as the threshold to screen significantly different phosphorylation state change.

### CRISPR/Cas9 knockout of CHS2 gene in susceptible strain JC-S

A pair of sgRNAs against exon 2 and exon 3 of *CHS2* gene was designed and synthesized as described above. After injection of the mixture of dual sgRNAs and Cas9, genomic DNA from exuviates were extracted using the Multisource Genomic DNA Miniprep Kit. Detecting primers were designed flanking the CRISPR target sites (Table S7). Three steps were taken to obtain hereditary homozygote: (i) reciprocal crossed between moths of G_0_ and wild type moth in single pair, and the mutation type of G_0_ moth was determined after laying enough eggs; (ii) G_1_ larvae of selected mutation type were reared to pupa, and determined it’s genotype; individuals carried the mutated allele were mass-crossed to produce G_2_; (iii) G_2_ individuals were genotyped, and homozygotes were selected to produce a knockout strain, named CHS2-KO. Diet overlay bioassay was used to test first instar larvae of CHS2-KO against Vip3Aa as described above.

### Fluorescence in situ hybridization (FISH) of *CHS2*

Midgut of fifth instar larvae from JC-S and CHS2-KO were used for RNA FISH analysis. RNA-FISH assays were performed according to the manufacture’s protocol provided by Stellaris FISH Probes (RiboBio) as described ^73^. Briefly, midgut section were permeabilized in 1 × PBS containing 0.5% Triton X-100 for 5 min at 4 ℃, then washed in 1 × PBS for 5 min. 200 μL of Pre-hybridization buffer was added at 37 ℃ for 30 min. Hybridization was carried out with a FISH probe in a moist chamber at 37 ℃ away from light more than 10h using Ribo Fluorescent In Situ Hybridization Kit (RiboBio). The slides were washed three times with Wash Buffer Ⅰ (4 × SSC with 0.1% Tween-20), once each with Wash Buffer Ⅱ (2 × SSC), Wash Buffer Ⅲ (1 × SSC) at 42 ℃ in the dark for 5 min and once with 1 × PBS at room temperature. Then, slides were stained with DAPI in the dark for 10 min. FISH probes of CHS2 was designed and synthesized by RiboBio Co., Ltd. Images were obtained with a confocal microscope (Nikon).

## Acknowledgement

This project was funded by the STI 2030 - Major Projects (2022ZD04021), Innovation Program of Chinese Academy of Agricultural Science (CAAS-CSCB-202303) and The Agricultural Science and Technology Innovation Program (CAAS-ZDRW202412).

## Data availability

The genome of JC-S and JC-R were submitted to Figshare database (10.6084/m9.figshare.26309923).

## Author Contributions

Conceptualization, Y.X., and M.J; Methodology, M.J., Y.S., Y.P., S.C., and X.Z; data analyze, M.J. and Y.P.; writing-original draft, M.J., writing-review & editing, M.J., K.L., and Y.X.; visualization, M.J. and Y.P.

