## supplemental files for "CRISPR-mediated chromosome deletion facilitates genetic mapping of Vip3Aa resistance gene within complex genomic region in an invasive global pest"

**Table S1. Responses of strain JC-R (selected with Vip3Aa) and its parent susceptible strain JC-S of *S. frugiperda* to Vip3Aa, Cry1Ab, Cry1Fa, and Cry2Ab.**

| Toxin | Strain | n[a] | $LC_{50}$[b] (95% FL[c]) | Slope (SE)[d] | RR[e] |
|---|---|---|---|---|---|
| Vip3Aa | JC-S | 336 | 0.048 (0.040-0.057) | 2.2 (0.2) | 1.0 |
| | JC-R | 336 | 250.35 (203.29-316.28) | 1.9(0.2) | 5247.6[f] |
| Cry1Ab | JC-S | 144 | 0.20 (0.15-0.26) | 2.3 (0.3) | 1.0 |
| | JC-R | 144 | 0.22 (0.17-0.28) | 2.6 (0.4) | 1.1 |
| Cry1Fa | JC-S | 144 | 0.015 (0.011-0.019) | 2.63 (0.4) | 1.0 |
| | JC-R | 144 | 0.020 (0.015-0.028) | 2.0 (0.3) | 1.3 |
| Cry2Ab | JC-S | 144 | 0.14 (0.11-0.19) | 2.1 (0.3) | 1.0 |
| | JC-R | 144 | 0.13 (0.10-0.17) | 2.4 (0.3) | 0.9 |

[a] Number of larvae tested.

[b] Concentration (μg toxin per $cm^2$ diet) killing 50% of larvae.

[c] 95% fiducial limits.

[d] Slope of the concentration-mortality line and its standard error.

[e] Resistance ratio = $LC_{50}$ for JC-R divided by $LC_{50}$ for JC-S for the same toxin.

[f] $LC_{50}$ of Vip3Aa significantly greater for JC-R than JC-S based on non-overlap of 95% FL.

**Table S2. Survival of first instar of *S frugiperda* strains JC-S and JC-R on Vip3Aa corn and non-Bt corn.**

| Strains | Non-Bt corn | | Vip3Aa corn | | Control efficiency of Vip3Aa corn[a] |
|---|---|---|---|---|---|
| | Total number | Survival number | Total number | Survival number | |
| JC-S_1 | 10 | 10 | 10 | 0 | |
| JC-S_2 | 10 | 9 | 10 | 0 | 93.3% |
| JC-S_3 | 10 | 9 | 10 | 0 | |
| JC-R_1 | 10 | 9 | 10 | 9 | |
| JC-R_2 | 10 | 10 | 10 | 8 | 6.7% |
| JC-R_3 | 10 | 9 | 10 | 9 | |

[a] Control efficiency was calculated: (average death number of strain feeding on Vip3Aa corn – average death number of strain feeding on non-Bt corn)/total number*100%.

**Table S3. Larval survival of S. frugiperda strains JC-R, JC-S, and their F1 progeny on diet treated with 0.4 µg Vip3Aa per cm$^2$ diet.**

| Strain or cross[a] | Survival (%)[b] | Dominance (*h*)[c] |
|---|---|---|
| JC-S | 0 | |
| JC-R | 100 | |
| JC-S ♀ vs JC-R ♂ | 6.3 | 0.064 |
| JC-S ♂ vs JC-R ♀ | 8.3 | 0.085 |

[a] F1 progeny were obtained from reciprocal mass crosses between JC-S and JC-R.

[b] Sample sizes were 48 larvae per strain.

[c] *h* = (Survival of F1 progeny - survival of JC-S)/(Survival of JC-R - survival of JC-S); *h* = 0 indicates completely recessive resistance, *h* = 1 indicates completely dominant resistance.

**Table S4. Fine-scale mapping using BC populations: seven markers on chromosome 18 tightly linked with resistance to Vip3Aa in strain JC-R of *S. frugiperda*.**

| Marker | Site (Mb) | *rr*[a] | *rs*[a] | $\chi^2$ | *P*[b] |
|---|---|---|---|---|---|
| 1 | 5.86 | 86 | 10 | 62.67 | 2.44E-15 |
| 2 | 6.93 | 96 | 0 | 100 | 1.52E-23 |
| 3 | 7.75 | 96 | 0 | 100 | 1.52E-23 |
| 4 | 8.51 | 96 | 0 | 100 | 1.52E-23 |
| 5 | 9.48 | 96 | 0 | 100 | 1.52E-23 |
| 6 | 10.46 | 96 | 0 | 100 | 1.52E-23 |
| 7 | 10.70 | 88 | 8 | 69.44 | 7.86E-17 |

[a] *rr*, homozygous for the marker from JC-R; *rs*, heterozygous from JC-R and JC-S. Based on analysis of 96 larvae that were backcross progeny and survived exposure to diet with 5.0 µg Vip3Aa per $cm^2$ diet.

[b] Probability based on the null hypothesis of random assortment *rr*:*rs* = 1:1.

**Table S5. Fine-scale mapping using $F_6$ populations: seven markers on chromosome 18 tightly linked with resistance to Vip3Aa in strain JC-R of *S. frugiperda*.**

| Marker | Site (Mb) | *rr*[aa] | *rs*[aa] | *ss*[aa] | $\chi^2$ | *P*[b] |
|---|---|---|---|---|---|---|
| 1 | 6.93 | 90 | 6 | 0 | 252.34 | 2.49E-53 |
| 2 | 7.52 | 96 | 0 | 0 | 300 | 2.90E-63 |
| 3 | 7.75 | 96 | 0 | 0 | 300 | 2.90E-63 |
| 4 | 8.51 | 96 | 0 | 0 | 300 | 2.90E-63 |
| 5 | 9.48 | 96 | 0 | 0 | 300 | 2.90E-63 |
| 6 | 10.06 | 96 | 0 | 0 | 300 | 2.90E-63 |
| 7 | 10.46 | 92 | 3 | 1 | 267.60 | 1.64E-56 |

[a] *rr*, homozygous for the marker from JC-R; *ss*, homozygous for the marker from JC-S; *rs*, heterozygous. Based on analysis of 96 larvae that were $F_6$ progeny and survived exposure to diet with 5.0 μg Vip3Aa per $cm^2$ diet.

[b] Probability based on the null hypothesis of random assortment *rr*:*rs*:*ss* = 1:2:1.

**Table S6. Statistics of JC-S and JC-R genome assemblies.**

| **Feature of genome assembly** | **JC-S** | **JC-R** |
|---|---|---|
| Total length of assembly (Mb) | 408.7 | 383.4 |
| Total number of scaffold | 140 | 77 |
| Scaffold N50 (Mb) | 13.3 | 13.4 |
| Longest scaffold (Mb) | 22.2 | 22.3 |
| Completer BUSCOs | 99.3% | 99.2% |

**Table S7. Primers used in this study.**

| Primers | Sequences | Function |
|---|---|---|
| Marker_1F | CTGGAATTATACGCTTTGTTGGGA | Fine-mapping |
| Marker_1R | CAATTTTCACTGTTGACTCGA | |
| Marker_2F | TTGACGTAGTTAACTAATAAGT | |
| Marker_2R | ATTATTTGTGTTGATGTAGTT | |
| Marker_3F | TGATAAATAAGAAGGTATTCG | |
| Marker_3R | CAAATTCATCGCTCGATACC | |
| Marker_4F | GAATAGTACTAATTTTGTAA | |
| Marker_4R | CTAGGAATAGTATTCGATAT | |
| Marker_5F | TGTGGGTGGTGCTGATCCAGG | |
| Marker_5R | GGGATCTGCGTTCCTCTCGC | |
| Marker_6F | CATTATTGAACTGTACCACCC | |
| Marker_6R | CGACTACACAAACTTATGAA | |
| Marker_7F | TACGCTTCTAATGTTAAGGTTC | |
| Marker_7R | GATAAAACAGTAATCTGGATG | |
| Marker_8F | TAGATATTTACGTACGACGGCGA | |
| Marker_8R | TGCAAAGTTGCGCAAGCCAG | |
| Marker_9F | GTTATACATGTACACGTAATA | |
| Marker_9R | CAAATACACGTTCTTTTATACT | |
| sgRNA1 | AATATTGTGCAAGATGCTCGTGG | sgRNAs for Knockout |
| sgRNA2 | TTCCGAATAAAAACTGCAACCGG | |
| sgRNA3 | ATTCCTAGGATTAGCATCATGGG | |
| sgRNA4 | GTGAACCTCTCCGTACTGGCTGG | |
| sgRNA5 | TCAGGCGACGAAGAAATGGATGG | |
| sgRNA6 | GGCACATACCTAGAGGAGCTGGG | |
| sgRNA7 | TCTCCAAAAGTTGTGGTCTTTGG | |
| sgRNA8 | GTGCCGCTAACATGCTGCTAATGG | |
| sgRNA9 | GAAAGGCCTGCTGATACGTCCCGG | |
| sgRNA10 | GCAGCCAACTATAGATGTTCAGG | |
| sgRNA11 | GAAGTCGCTAAGATAGTGGCCGG | |
| Detect_1F | GTACACGTTTGTCGCGTCCCGAACACG | Mutation detecting of knockout |
| Detect_1R | ATTTATTCGCAGTTTATGGCCTCT | |
| Detect_2F | TATAAACTGAGAAAACAAACGAA | |
| Detect_2R | GTCAGTAAGGCTGCCAACAAGG | |
| Detect_3F | ATATAGCAATGAGAGATTCT | |
| Detect_3R | ACTGGGATCCACAGCGATAGC | |
| Detect_4F | GCTCCAGGCCGGCGTGCGCCC | |
| Detect_4R | GCGTGACATGGCCGGCCACCGG | |
| Detect_5F | CACACATGTGTCCTCGCTGCCAG | |
| Detect_5R | GTGTCGGTCCGATGAGGGTCGG | |
| Detect_6F | AGTAAGCTGTAGGCGATCAACAAA | |
| Detect_6R | GCGGCGCAGGCGAAGCAGCGC | |
| Detect_7F | AAGGTGTCGGACCGCACGTGGC | |

| | | |
|---|---|---|
| Detect_7R | GCCGGTCTGGGTTTCCACTG | |
| Detect_8F | TCGTGAGGATGGTTTCTGTGAT | |
| Detect_8R | TCTTAGCATCACTTTTCTT | |
| Detect_9F | AAGTTCCTTCTAGAGTAGCAGATA | |
| Detect_9R | ACTGCATTCTGCACACTTCCAA | |
| Detect_10F | TAGTTGTGTGGGACAATGGAACT | |
| Detect_10R | GATTACGTAAACGCAATCCC | |
| sgRNA1-CHS2 | GGATCTGCGGTTGTGTCTAA | sgRNAs for CHS2 knockout |
| sgRNA2-CHS2 | GCATTCAGTATTCCAGAAGT | |
| Detect-CHS2-F | AGGATGGAATCTGTTTCGAGAGA | Mutation detecting of CHS2 knockout |
| Detect-CHS2-R | AGAAGGCTTCGGTGCTGTT | |

**Supplementary Figures**

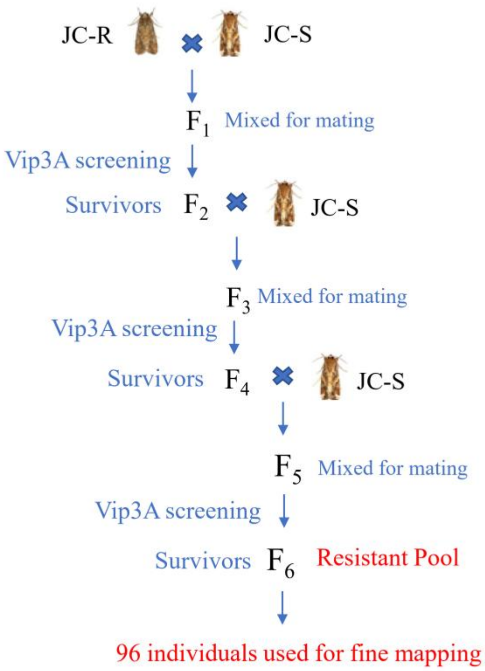


**Figure S1. Construction of a $F_6$ population for fine-mapping.** For fine mapping, a single-pair cross was conducted between JC-R (female) and JC-S (male). The $F_1$ progeny were raised on a normal diet, resulting in the production of $F_2$ progeny. Then, approximate 1000 $F_2$ larvae were screened with Vip3Aa containing diet (5.0 µg Vip3Aa per $cm^2$ diet). The survivors were reared to adult and crossed with JC-S to produce $F_3$. A total of six generations was generated. Ninety-six survivors in $F_6$ generation were selected for fine mapping.

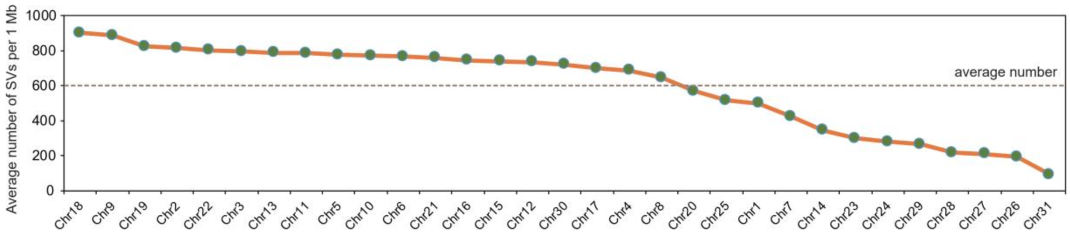


**Figure S2. The average number of SVs per 1Mb in each chromosome.**

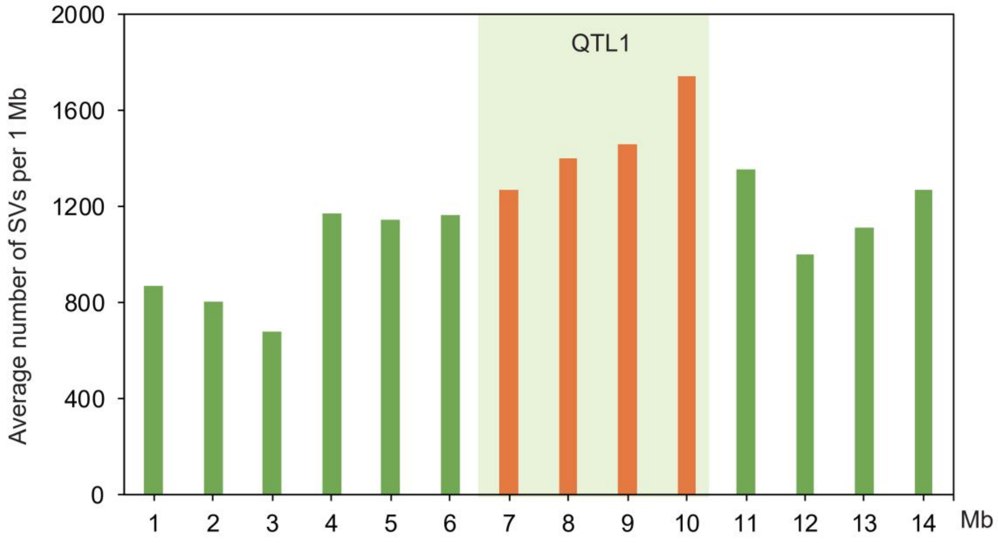


**Figure S3. The average number of SVs per 1Mb in chromosome 18.** LOC1 region was labeled in red.

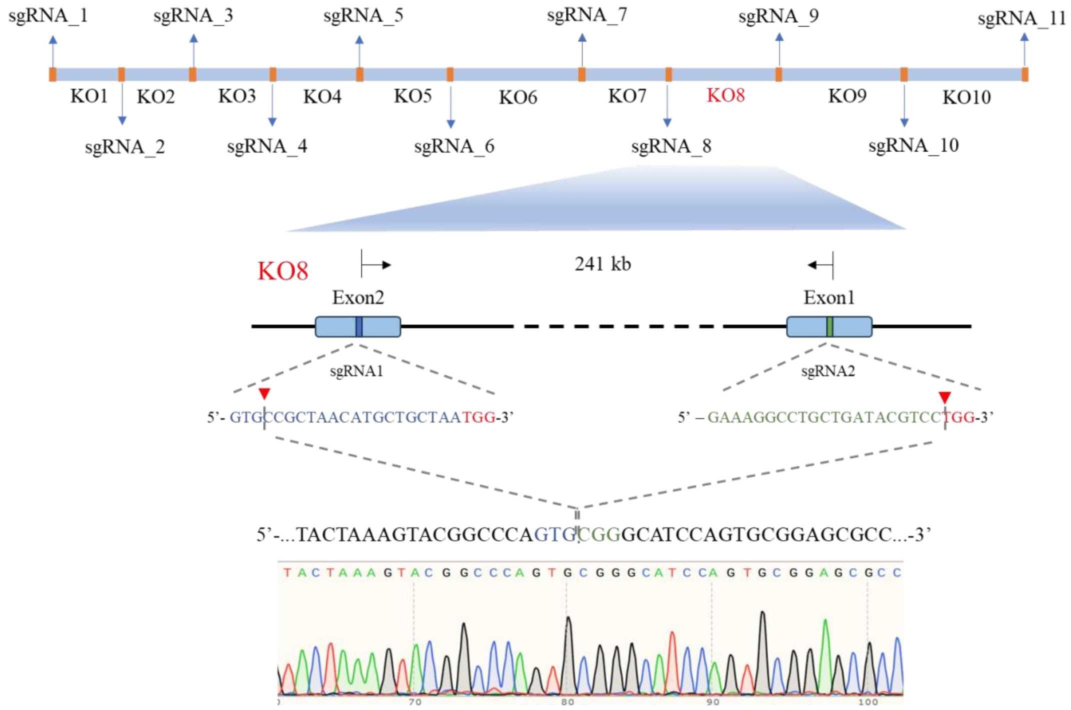


**Figure S4. sgRNAs designed for chromosome fragment stepwise knockout.** A total of 11 sgRNAs were designed. Two adjacent sgRNAs were combined to knockout chromosome fragment. The detail information of the KO8 region which containing the resistant gene. The KO8 region was deleted 241 Kb yielding a chromosome deletion. sgRNA target sequences (blue and green), protospacer adjacen motif (PAM, red), Cas9 cleavage site (red triangle). Chromatograms of direct sequencing of PCR product for determining genotype.

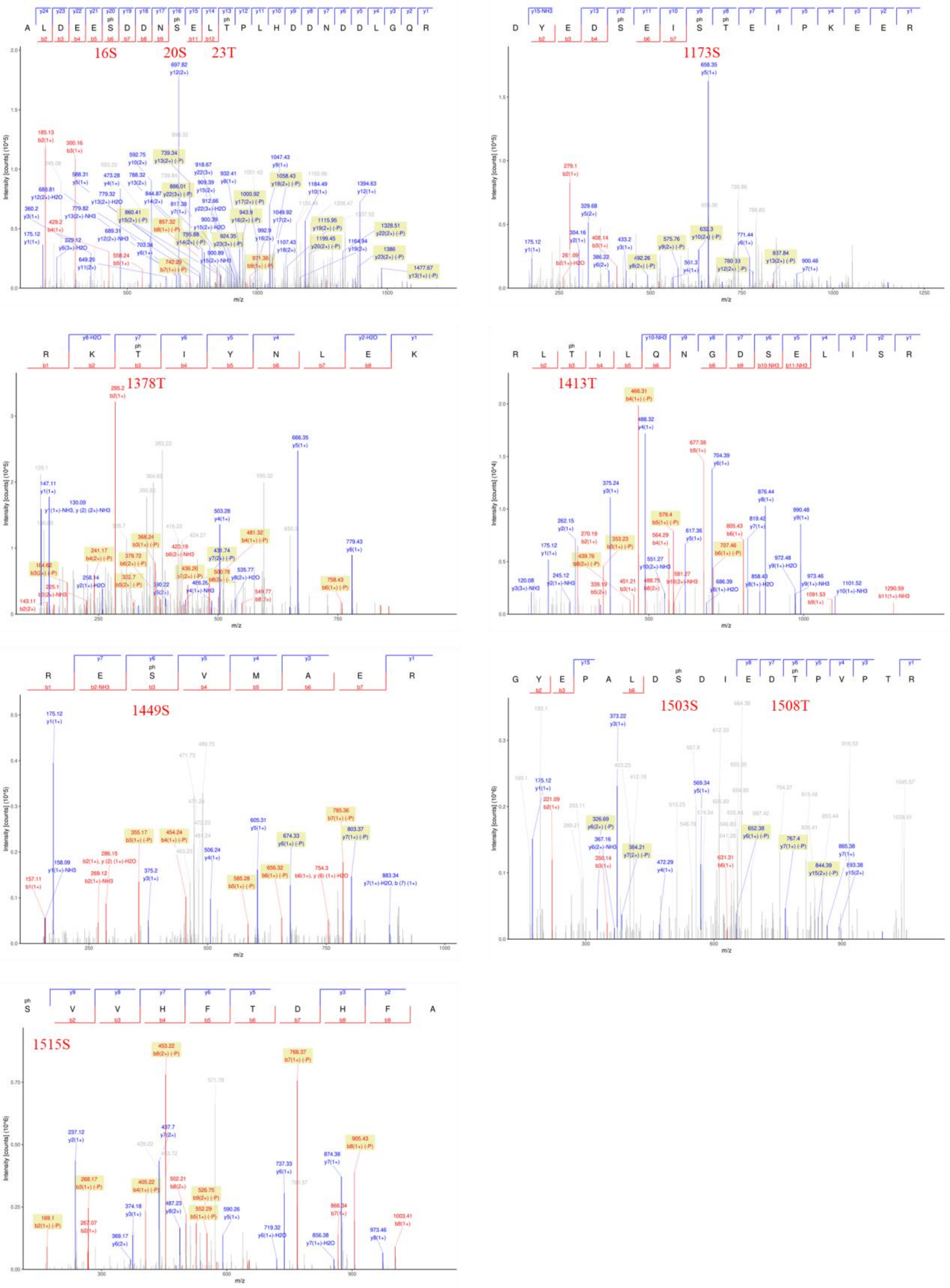


**Figure S5. The putative phosphorylation sites of pS16, pS20, pT23, pS1173, pT1378, pT1413, pS1449, pS1503, pT1508, and pS1515 of CHS2 were identified by LC-MS/MS.** The identified phosphorylation sites are marked in red.

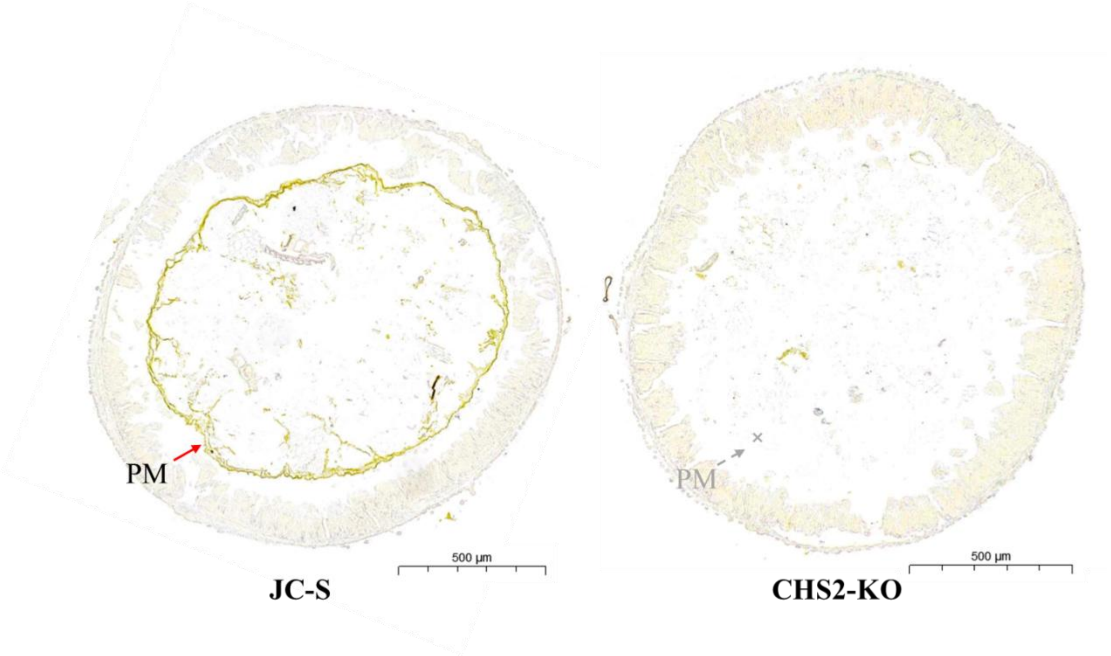


**Figure S6. The colons of midgut from JC-S and CHS2-KO strain.** In JC-S, the PM is clearly visible. However, the PM was disappeared in the CH2-KO strain.
